# Correlated traits generate the structure and stability of multiplex ecological networks

**DOI:** 10.64898/2026.09.30.755657

**Authors:** Kayla R. S. Hale, Elisa Thébault, Fernanda S. Valdovinos

**Affiliations:** Santa Fe Institute; Santa Fe, New Mexico, USA 87501; Centre for Ecosystem Management, Department of Integrative Biology, University of Guelph; Guelph, Canada N1G 2W1; Institute of Ecology and Environmental Sciences, Campus Pierre et Marie Curie, Sorbonne Université, CNRS; Paris, France 75006; Department of Environmental Science and Policy, University of California; Davis, California, USA 95616

**Keywords:** Biological sciences, ecology, network topology, species interactions, community matrices, multiplex networks

## Abstract

Ecological stability and function emerge from the many and diverse types of interactions among species. Ecological networks represent the structure and strength of species interactions, but their study is often restricted to single interaction-types, likely due to an absence of models for plausibly (i.e., nonrandomly) structured ‘multiplex networks’ of multiple interaction types. Here, we present a general but mechanistic model that simulates multiplex networks using the different, but correlated, traits that determine different species interactions. We show that ecological stability varies systematically with these trait correlations by controlling the emergent complexity and positioning of strong interactions in the community. Our model provides a flexible but power tool to study multiplex networks under alternative trait-based hypotheses, offering new insights into the structure and stability of complex natural systems.

**One-Sentence Summary:** Trait correlations structure stability in complex ecological communities of multiple interaction types.

**Significance Statement:** To effectively tackle the increasing impacts of global change on ecological systems, we need to understand how species interactions may propagate or buffer disturbance, and the mechanisms by which resilience to disturbances emerge. This topic has been investigated for random communities and single interaction-type (e.g., predator-prey) networks. Yet, natural systems include many diverse types of interactions, and feedbacks between them mediate how species and ecosystems respond to global change. Here, we present a general model to study ‘multiplex networks’ of multiple interaction types generated from ecological traits. We show that resilience varies systematically with trait correlations underlying the structure and strength of interactions involving key species that interconnect, and therefore propagate disturbances between, subnetworks of different types of ecological interactions.

## Introduction

Ecological networks provide powerful conceptual and technical tool for analyzing the complexity of species-rich ecosystems. Studies of network structure have shown that species interactions are organized into predictable, nonrandom patterns determined by species’ traits^1–10^. These patterns in turn shape ecosystem respond to perturbations. For instance^11–15^, the effects of extinctions on biodiversity and ecosystem function strongly depend on the traits, position, and partners (the *functional roles*), of removed species. However, documenting empirical networks is costly, both in terms of time and expertise^10,16,17^. Stochastic generating models help fill this gap by producing plausible, trait-based network replicates with which to study the relationship between structure and stability in complex ecological systems^9,10^.

Yet, these models have so far been restricted to networks composed of single interaction at types, i.e., *subnetworks* of the broader community^16,18–21^. Natural systems include diverse types of interactions among co-occurring species, which can be represented as *multiplex networks* of subnetworks interconnected by species that participate in multiple interaction types^16,22^. The functional roles of these key species can strongly affect ecosystem stability and function^18,23–25^ by propagating perturbations between different subnetworks. For instance, herbivores affect the structure and dynamics of mutualist subnetworks through their impacts on plants^26–28^, and these changes can indirectly propagate to alter landcover and the behavior of top predators^29^.

Understanding how feedbacks among different interaction types shape natural systems has therefore been a goal of ecological research for more than 20 years^9,12,16,18–20,22,30–34^. Though empirical data have begun to be documented^21,35–45^, they often exclude important interconnections among subnetworks, artificially imposing ‘multipartite’ structure. At the same time, theoretical studies are largely restricted to random communities^46–51^ or randomized versions of empirical networks^24,52–54^ (but see^55^).

Here, we present a general, trait-based model for multiplex ecological networks. We find that trait correlations underlying the interconnection of subnetworks into multiplex networks can strongly impact stability by altering emergent community complexity, and independent of complexity, by controlling the relative positioning of the strongest interactions in the community.

### The multiplex model

Stochastic generating models^2,3,8,56–61^ have been an essential tool for advancing ecological and evolutionary research^9,10^ by allowing researchers to test how ecological mechanisms shape network structure^2,3,8,56–61^ and, ultimately, community stability^46,47,62,63^. In these models, species are randomly assigned traits from a probability distribution (ordering them along a *trait axis*), and *interaction rules* determine which pairs of species can interact based on their trait values. This process is repeated stochastically to generate a set of simulated networks that can be compared to empirical networks as null hypotheses for emergent structure (e.g.,^64^), or used as replicates in theoretical studies where empirical data remain limited (e.g.,^65^).

Simple rules derived from classic stochastic generating models have been shown to successfully replicate many features of empirical subnetworks^2,9,58,66–68^, especially compared to random networks. For example, pollinators tend to access plants below a threshold of flower corolla length matching the length of their mouthparts (*threshold rule*^58,68^), whereas predators tend to hunt prey within a contiguous range (i.e., an interval) of body sizes smaller than their own (*interval rule*^60,68^). The success of these models reflects, in large part, that empirical networks of a single interaction type are low dimensional, meaning that they can typically be explained by a single or at most a few trait axes^4,69^. Although these axes may be abstract, they are typically grounded in measurable ecological traits^4,70^. Further, different interaction types tend to be mediated by different traits^4,10,70^, such as body size for predator-prey interactions^69^, flower and mouthpart traits for pollination^71^, and physical and chemical defenses for herbivory^72^.

Critically, these traits are unlikely to be independent. Shared ecologies and evolutionary histories^27,32,73–77^, including selection pressures from natural enemies, competitors, and mutualists, as well as organismal allometries, genetic correlations, and ecological tradeoffs in performance, shape multiple traits simultaneously. These processes generate correlations among traits that should also shape how species function across multiple interaction types^24,32,77^. For example, the same phenolic compounds that attract pollinators and seed dispersers to flowers and fruits also attract herbivores and seed predators, such that plants supporting the most mutualists may also be the most vulnerable to natural enemies^24,32,73^.

We build on this framework to introduce a stochastic generating model for multiplex ecological networks – subnetworks interconnected by shared species – using correlated trait axes (**Fig. 1**; see Methods). Rather than assigning traits independently for each subnetwork, we model species’ traits as jointly distributed using a *copula*, a multivariate probability distribution specifying the joint trait structure for the community. When species participate in multiple interaction types, underlying trait correlations shape their functional roles across subnetworks, and therefore emergent multiplex structure.

**Figure 1.**
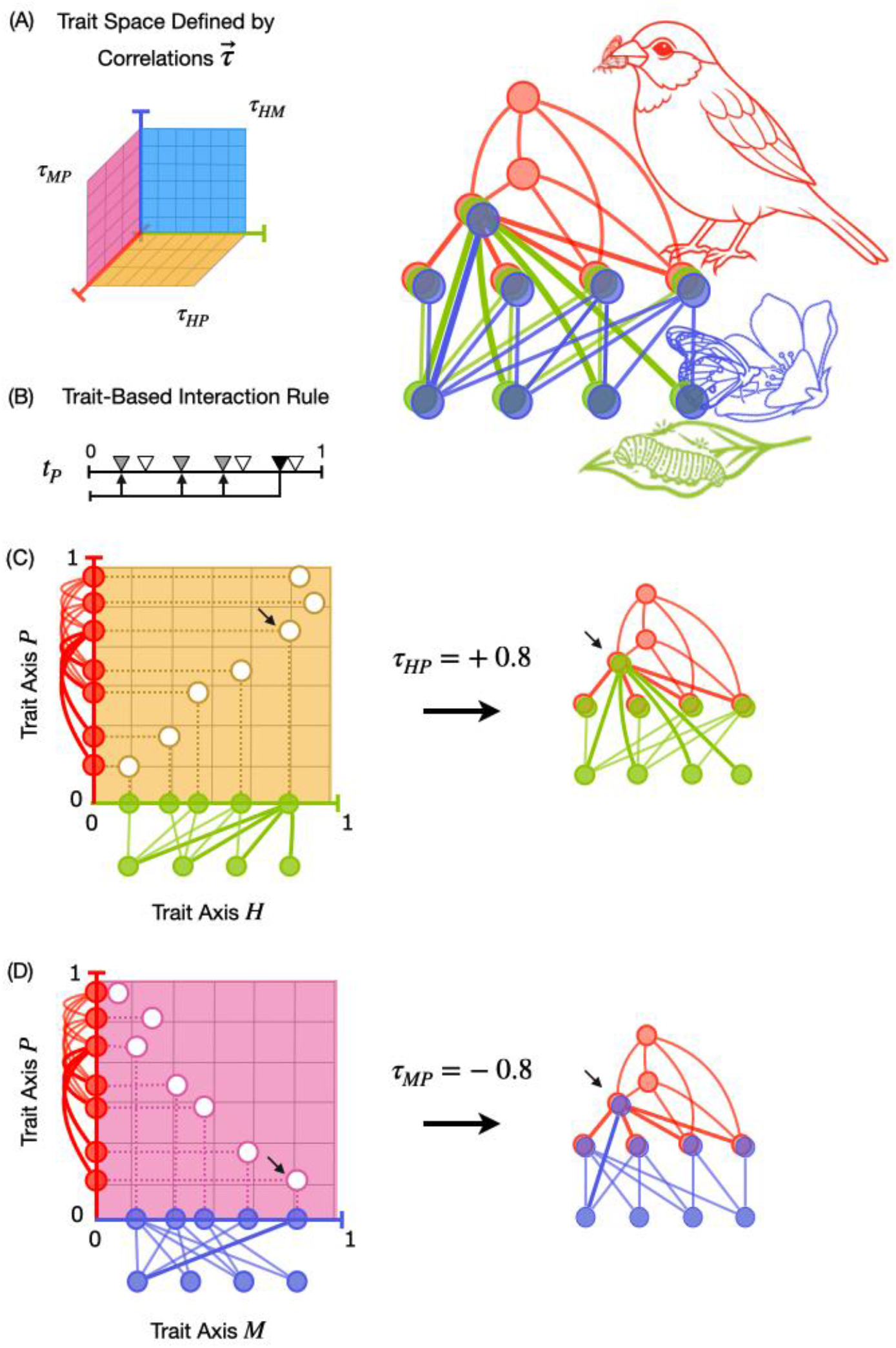
Multiplex networks are constructed from subnetworks of different interaction types generated by correlated trait axes. We illustrate our model for a community of *S* = 11 species assembled into a multiplex network (**upper right**) with predator-prey (red), animal-plant mutualism (purple), and animal-plant herbivory (green) interactions determined by trait *x* ∈ {*P, M, H*}, respectively, and their pairwise trait correlations *τ*_*MP*_, *τ*_*HP*_, and *τ*_*HM*_ (pink, yellow, and blue, respectively). Traits for each species are randomly sampled from a copula such that traits *x, y* have specified correlation *τ*_*xy*_ while remaining uniformly distributed along each trait axis (**A**, 3-D space from which the copula is sampled). An interaction rule (**B**) is then applied independently to each trait axis to generate the corresponding subnetwork. Here, we use the threshold rule (^68^, derived from ^58^), in which a species can potentially interact with any species below its own (threshold) trait value. The multiplex network (**upper right**) is simply the set of subnetworks interconnected by species that participate in multiple interaction types. Trait correlations therefore do not affect the structure of the subnetworks but rather species’ positions and partners (i.e., their functional roles) across subnetworks. For example, in (**C**), the strong positive correlation between trait axes *H* and *P* causes species with high trait values, and therefore a high number of partners, in one subnetwork to also tend to have high number of partners in the other (e.g., species marked with arrow). In (**D**), the strong negative correlation between trait axes *M* and *P* reverses this relationship while preserving the structure of the individual subnetworks. In this illustration (following a simplified version of ^64^), predators (e.g., birds) can prey on both herbivores and mutualists, both herbivores and mutualists forage on plants, and some animals even act as both herbivores and mutualists simultaneously or at different lifestages (e.g., larval caterpillars [herbivores] transition to adult butterflies [pollinators]). Image credits: Caterpillar Icon #5544091, Icongeek26; Leaf Icon #5003436, Przemyslawk; Monarch Icon #5929076, Theodoros Lamprou; Sparrow Icon #6594577, Fauzi Arts; all images licensed by The Noun Project.

Our model generates a multiplex network of *S* species consisting of *n* subnetworks, as follows. For each subnetwork, we define a trait axis *x* and an interaction rule (**Fig. 1B**). Each species *i* ∈ {1, …, *S*} is assigned a set of traits 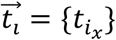 drawn from an *n*-dimensional Gaussian copula. Values for each trait *t*_*x*_ are uniformly distributed along their axis (marginal distributions) but are constrained by correlations between each pair of traits *τ*_*xy*_ (**Fig. 1C-D**). Then, we randomly sample *S*_*x*_ species from each trait axis and apply the corresponding interaction rule to achieve a simulated subnetwork with a specified connectance (*C*_*x*_, the fraction of possible interactions that are realized). The multiplex network emerges from species participating in multiple subnetworks. As the species richness of subnetworks *S*_*x*_ increase relative to overall multiplex species richness *S*, species become more likely to participate in multiple interaction types simultaneously.

The topological properties of each subnetwork are invariant to the underlying trait correlations, whereas multiplex properties, such as the composition and functional roles of species across subnetworks, vary systematically with the trait correlations (e.g., **Fig 1C-D**). Like classic generating models for ecological subnetworks, our multiplex model can be used as an ecologically informed null expectation for empirical multiplex networks. For example (Supplementary Text, *An empirical example from the Galapagos*), our model reproduces key properties of an empirical network^78,79^ wherein birds act as both seed dispersers and pollinators (“double mutualists”^80^) to their plant partners given the known correlation between two traits involved in these interactions (*τ*_*obs*_ ≈ 0.74 for beak widths and lengths^81^).

### Stability of multiplex communities

Here, we use our model to investigate, for the first time, how trait-driven multiplex network structure – that is, the pattern of interconnections among subnetworks of different interaction types – affects the stability of complex communities. Specifically, we study multiplex networks composed of three interconnected subnetworks (**Fig. 1, upper right**): unipartite predator-prey (antagonistic, indexed *P*); bipartite herbivore-plant (antagonistic; indexed *H*); and bipartite animal-plant mutualist (mutualistic; indexed *M*). These three subnetworks are foundational to natural ecosystems^17,64^, and though thought to be structured by different trait axes^4^, share similar generating mechanisms^70^ including trait matching^6^ and resource consumption^82^.

For each feasible combination of pairwise trait axis correlations (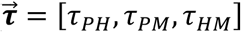, hereafter referred to as a network *treatment*), we generate a set of replicate multiplex networks using the threshold rule (see above) with fixed subnetwork richnesses (*S*_*x*_) and connectances (*C*_*x*_); this is the simplest rule that has been shown to reproduce key structural features in both food webs^58^ and mutualistic subnetworks^68^. Then, we convert each multiplex network replicate into a community matrix^83^ according to the following ecological assumptions (also see Methods): (i) *generalized Lotka-Volterra dynamics* linearized at a feasible equilibrium^84^, (ii) *additive interaction strengths* across subnetworks (i.e., multiple interaction types between two species have additive net effects^85^), (iii) *equally allocated foraging effort*, where the effects of consumers are split equally among all of their resources^48^, whether prey, plant, or mutualist, such that specialist consumers have stronger effects than generalists^47^, and (iv) *energetic inefficiencies* of consumption, such that consumers exert stronger effects on resources than the benefits they receive reciprocally (i.e., asymmetric interaction strengths^86^). Finally, we calculate the maximum real eigenvalue 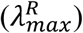 of the Jacobian of the community matrix to determine stability. A community is asymptotically locally stable, i.e., will return to equilibrium following a small perturbation, if 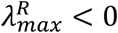. In this way, our study ties community trait structure to both the structure and strength of multiple interaction types in complex communities, and therefore to the emergent stability of multiplex ecological networks.

Community stability varies widely across our treatments (**Fig. 2A-C**), with the fraction of locally stable communities ranging from 12 to 71% depending on the underlying correlations between trait axes 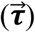. This represents a strong deviation from the 29% of stable communities when multiplex networks are constructed from uncorrelated trait axes 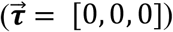, i.e., when subnetworks are interconnected randomly. Increasing the correlation between predator-prey and mutualist traits (*τ*_*PM*_), and to a lesser extent, the correlation between predator-prey and herbivore-plant traits (*τ*_*PH*_), tends to decrease community stability in our treatments, as quantified by their mean resilience 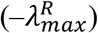 (**Fig. 2D**). Conversely, increasing the correlation between herbivore-plant and mutualist traits (*τ*_*HM*_) tends to increase resilience. To understand these effects, we distinguish between two consequences of trait-generated multiplex structure for community stability.

**Figure 2.**
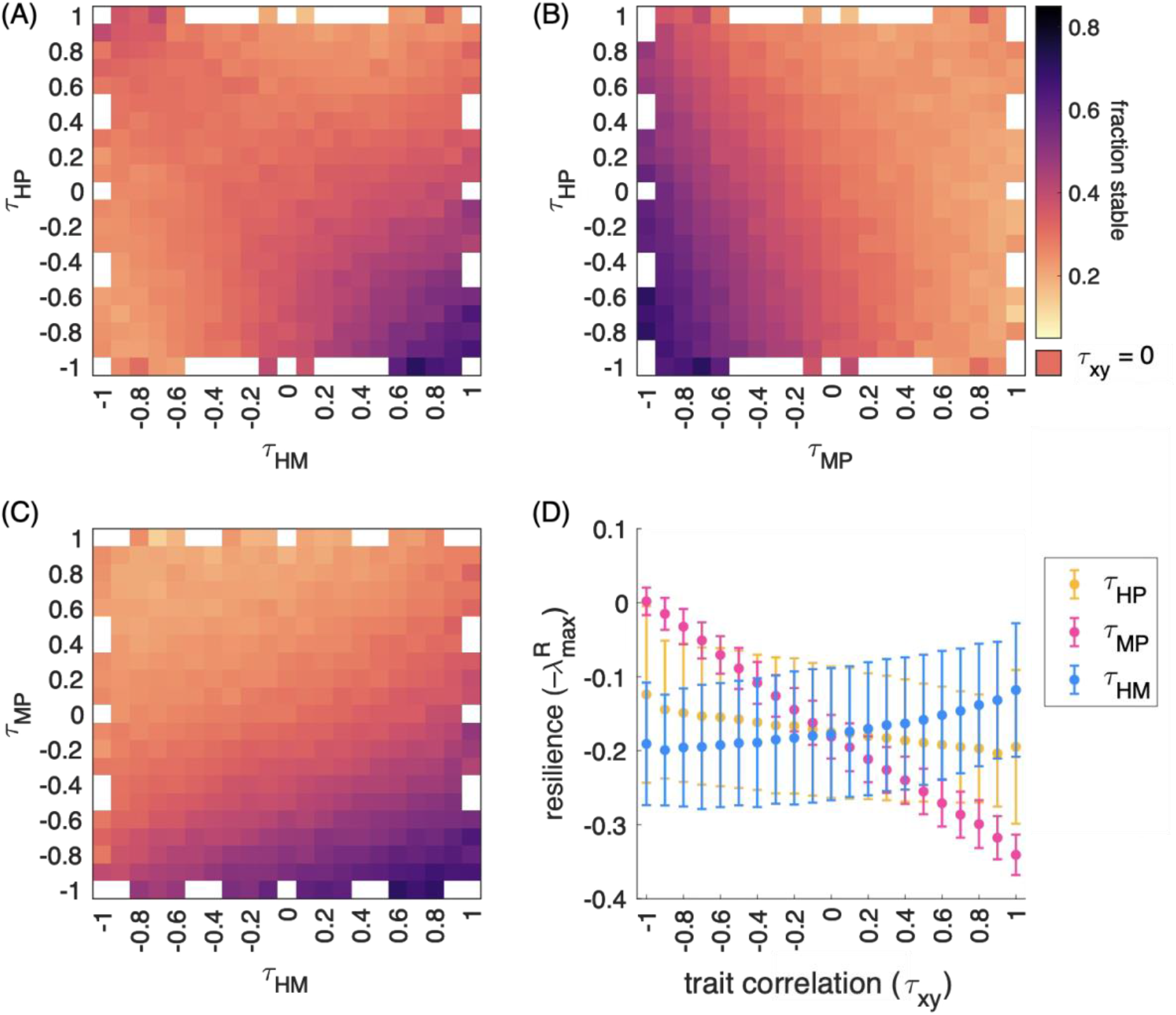
Community stability varies systematically with correlations among the traits underlying different interaction types. (**A-C**) Heatmaps show the effect of pairwise correlations (*τ*_*xy*_) among predator-prey (*P*), herbivore-plant (*H*), and animal-plant mutualist (*M*) trait axes on the community stability of multiplex networks. Multiplex networks were simulated in treatments of 300 stochastic replicates for each of the 4,787 sets of feasible trait correlations, 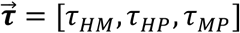. A replicate is locally stable if its resilience 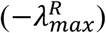 is positive, where 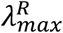 is the maximum real eigenvalue of the Jacobian of the corresponding community matrix. Heatmap cells show the fraction of locally stable replicates within a treatment, averaged over the set of treatments with a given set of pairwise trait correlation: (**A**) τ_HM_, τ_HP_, (**B**) τ_HP_, τ_HM_, and (**C**) τ_MP_, τ_HP_; darker colors indicate greater stability. Treatments generated from correlated traits ranged from 12% to 71% of locally stable replicates compared with 29% when trait axes were uncorrelated (i.e., when 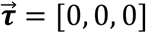). (**D**) Mean resilience decreases with a stronger correlation between predator-prey and herbivore-plant traits (τ_HP_, yellow) and between predator-prey and mutualist traits (τ_MP_, pink), but increases with a stronger correlation between herbivore-plant and mutualist traits (τ_HM_, blue). Points and error bars show the means and standard deviations among treatments with the indicated trait correlation *τ*_*xy*_. See **Supplementary Fig. S5** for stability results across all feasible sets of trait correlation combinations. Note that multiplex networks are of fixed complexity (species richness and number of links), but that community matrices can vary in emergent complexity due to species’ functional roles and the distribution of their foraging effort.

First, classic theory predicts that increasing community complexity 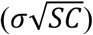, defined by greater diversity (*S*), connectance (*C*), and variation in interaction strengths (*σ*^2^), decreases stability in random and single interaction-type communities^46,50,63^. Trait correlations shape the emergent complexity of the community matrix (**Fig. 3A, Fig. S6**), with the associated changes in resilience (**Fig. 3B**) expected from this classic theory. Negative trait correlations (*τ*_*xy*_ < 0) tend to spread foraging effort more evenly across the community by aligning generalist consumers in one subnetwork with specialists in another, reducing the overall variance among interaction strengths (*σ*^2^) and increasing resilience. Positive trait correlations (*τ*_*xy*_ > 0) increase the probability of hyperspecialist consumers – species that exert the strongest effects by concentrating foraging effort on a single resource, whether prey or plant – thereby increasing *σ*^2^ and decreasing resilience. However, a strongly positive correlation between mutualist and herbivore traits (*τ*_*HM*_ → 1) is also associated with a higher probability that the same animal species are positioned as both mutualists and antagonistic herbivores to the same plants. By our assumption of additive interaction strengths, two distinct interaction links in the multiplex network are therefore represented as one weaker “dual” interaction in the corresponding community matrix (see^85,87^), decreasing overall community connectance (*C*) and increasing resilience.

**Figure 3.**
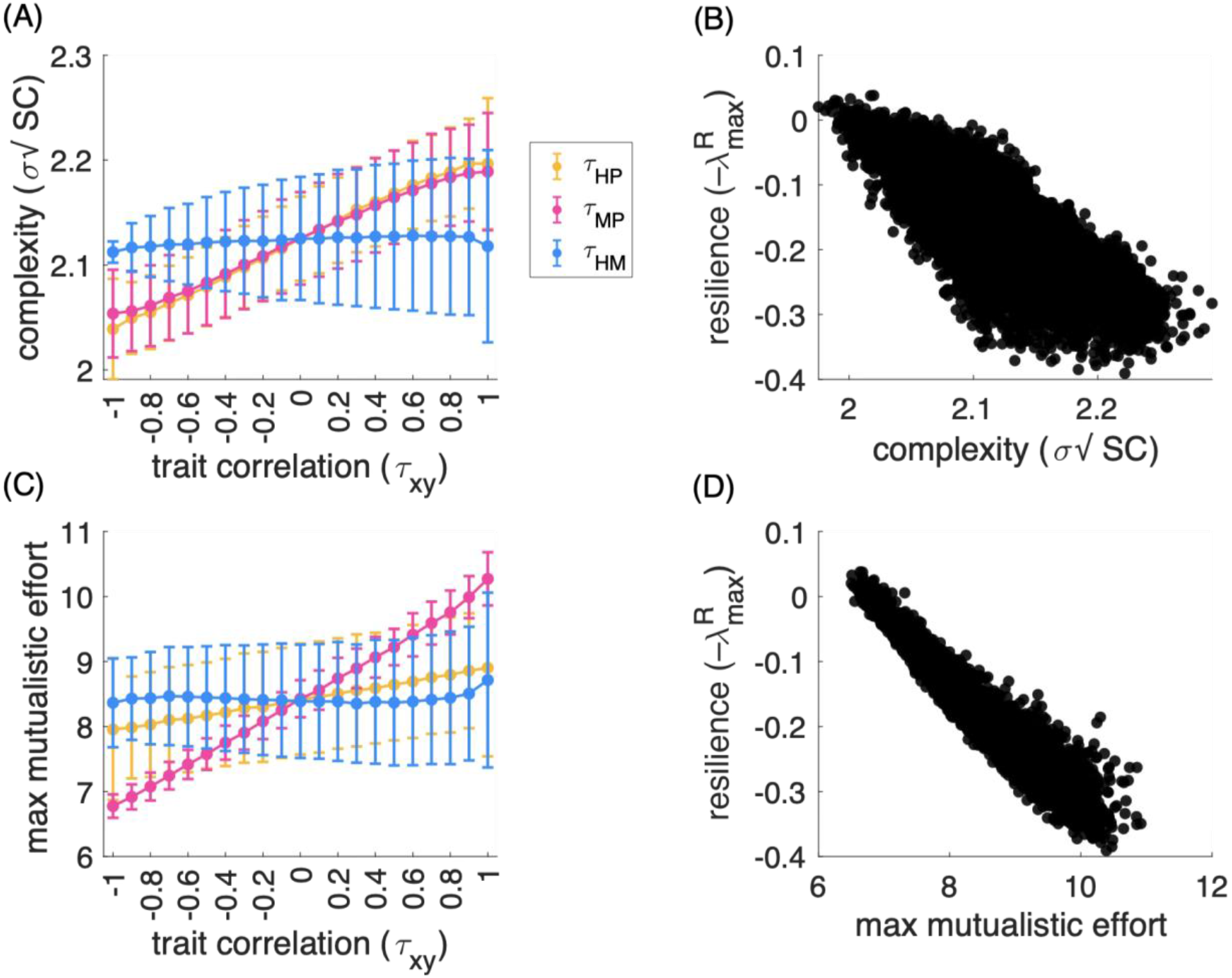
Trait correlations affect resilience through the emergent complexity and positioning of strong interactions in communities. **(A)** Trait correlations alter emergent community complexity through the distribution and alignment of consumers’ foraging effort across interaction types. Complexity 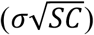 increases with the connectance (*C*) and variance of interaction strengths (*σ*^2^) in the community matrix (*C*), while species diversity (*S*) remains fixed. Negative correlations among the predator-prey and herbivore-plant (*τ*_*HP*_, yellow) and the predator-prey and mutualist-plant (*τ*_*MP*_, pink) trait axes align generalist consumers in one subnetwork with specialist consumers in the other, evenly distribution foraging effort across subnetworks and decreasing interaction strength variance (*σ*^2^) and community complexity as a whole; positive correlations have the opposite effect. Increasing the herbivore-plant and mutualist-plant trait correlation (*τ*_*HM*_, blue) aligns negative herbivory and positive mutualistic effects on shared plants, creating weak net effects that overall reduces effective community connectance (*C*), and therefore reducing complexity in such a way as to offset increases in (*σ*^2^) as *τ*_*HM*_ → 1. (**B**) Greater emergent complexity is associated with lower resilience among the multiplex network treatments. (**C**) Trait correlations also alter the relative concentration of foraging effort in the community. Increasing correlations between predator and mutualist traits (*τ*_*MP*_, pink) increases the frequency and co-occurrence of hyperspecialist mutualists that exert the strongest effects in the community. This concentrates mutualist foraging effort and strong positive interactions onto individual plant species. (**D**) Greater maximum mutualistic foraging effort concentrated onto a single species is associated with lower resilience. In (**A, C**) points and error bars show the means and standard deviations among treatment with the indicated trait correlation *τ*_*xy*_. In (**B, D**), each point represents the mean of one treatment (*N* = 4,787). See **Supplementary Fig. S6** for a breakdown of the results by each individual driver of complexity.

Second, beyond complexity, trait correlations affect resilience through the relative positioning of strong mutualisms within communities. The strong positive correlation between predator and mutualist traits (*τ*_*MP*_ → 1) increases both the probability of hyperspecialist mutualists and the frequency with which they concentrate mutualistic foraging effort onto the same plant resource (**Fig. 3C**). As maximum total concentration of mutualistic foraging effort within a community increases, resilience strongly decreases (**Fig. 3D**). These configurations create localized concentrations of the strongest interactions in the community, which may decrease stability by allowing positive feedback loops to become amplified through the same plant species.

To disentangle these effects from complexity, we use a representative subset of our multiplex network treatments to perform an additional experiment. For each treatment, we shuffle (i.e., randomize) the positioning of foraging effort within each of the replicate communities, while maintaining their complexity (*S, σ*, and *C*) and the reciprocal sign structure and strengths among species pairs (also see Methods). Interestingly, the shuffled communities are on average less resilient than their originals (**Fig. 4A**), indicating that the trait-generated structure of our multiplex networks confers greater stability than random communities with identical complexity and interaction strengths. Further, among these shuffled communities, greater concentrations of mutualistic foraging effort are associated with decreased resilience (**Fig. 4B**). Altogether, these results reveal that the stability of ecological communities is not determined solely by complexity, but also by the trait structure that shapes the strength and positioning of mutualistic and antagonistic interactions in ecological networks.

**Figure 4.**
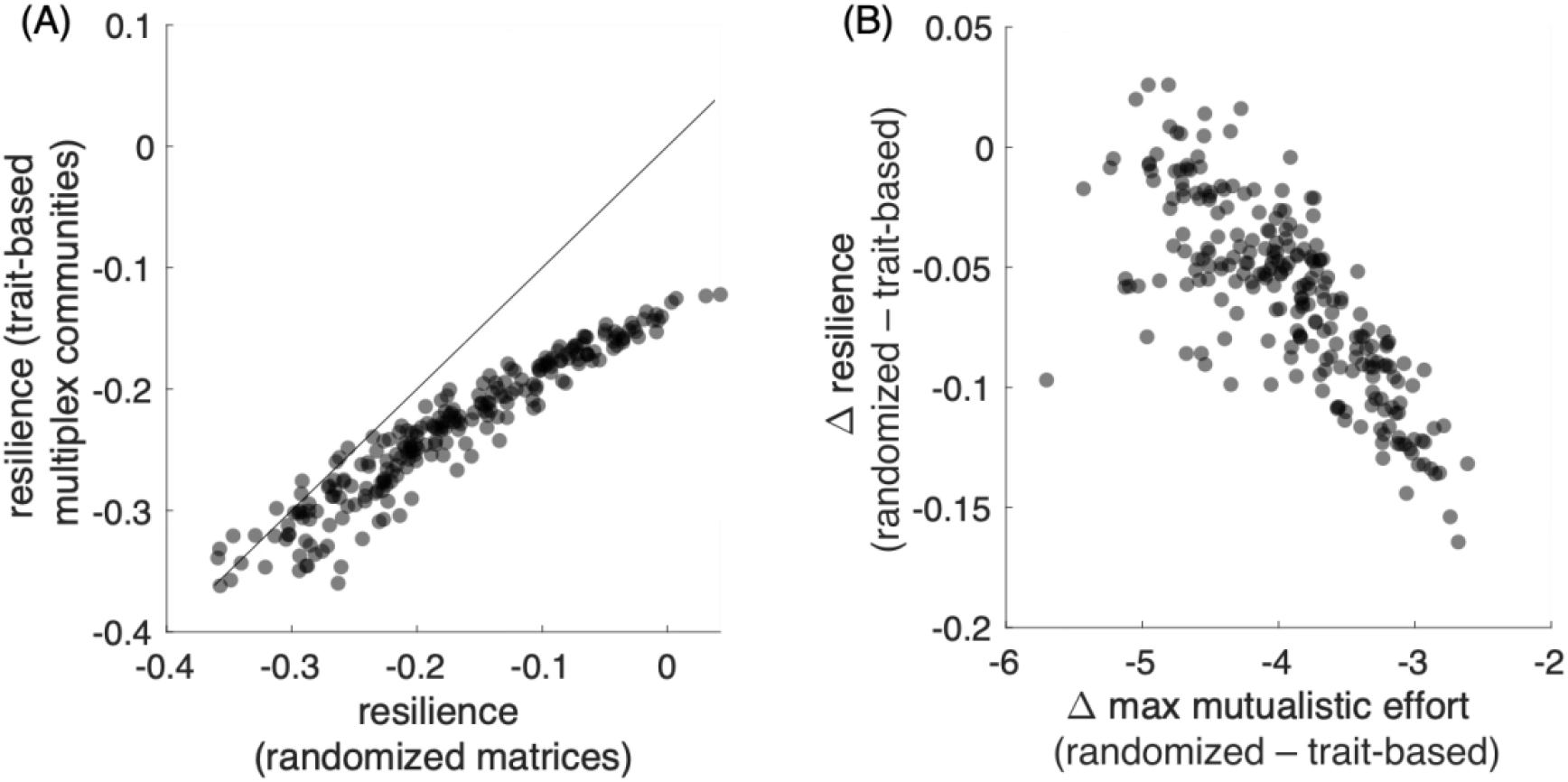
The strength and positioning of mutualistic interactions affect resilience independently of community complexity. For a subset of 240 multiplex network treatments spanning the trait correlation space 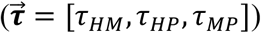, the positioning of consumers’ foraging effort within each trait-generated community matrix replicate was shuffled 100 (i.e., randomized) times while exactly preserving its complexity (connectance, mean and variance of interaction strengths, correlation and signs among reciprocal interaction strengths). (**A**) Randomized matrices were on average generally less resilient than their original trait-generated communities. The diagonal line shows the 1:1 expectation if randomization has no effect on resilience. (**B**) Increases in the maximum mutualistic foraging effort among randomized matrices with fixed complexity drive decreases in resilience, where changes (Δ) are presented as relative to the original trait-generated values (randomized – trait-based). (**A-B**) Each point is the mean value for one treatment (*N* = 240).

## Conclusions

For decades, ecologists have called for the synthetic study of species interactions in a general mechanistic framework^9,12,16,18–20,22,30–34^. Here, we introduced, for the first time, a tool that draws from the simple observations that species’ traits underly many different types of interactions, that these traits are unlikely to be independent, and further, that common mechanisms represented by simple rules can generate emergent network structures. Then, we demonstrated an application of this tool to studying the stability of complex communities structured by multiplex networks of three fundamental interaction types – predator-prey, plant-herbivore, and plant-animal mutualist.

Our results extend previous findings on the stability of random communities^46–51^, communities of single interaction types^62,86,88^, and specific empirical networks^24,52–54^. Correlations among species’ traits can significantly affect the functional roles (position and partners) of key species interlinking different subnetworks, and therefore their ability to propagate instability when perturbed. As a result, the effects of community structure on stability depend not only on the composition (i.e., relative proportions) of interaction types^46,47^, but also on their (nonrandom) structuring into multiplex networks, as determined by species’ interaction traits. Indeed, trait-generated structure tends to impart greater resilience to communities of multiple interaction types than expected from randomized versions of the communities with fixed complexity.

This work is only a first step towards understanding how the nonrandom structure of the many diverse types of ecological interactions in natural systems shape their stability and function. In the main text, for example, we used an interaction rule that has already been robustly studied^58,68^ to generate our subnetworks. However, the framework is general: any set of interaction rules based on trait axes can be used to generate subnetworks, including extant approaches not studied here^2,3,8,56–61^. In this way, our model provides a simple, fast, and customizable tool to encode and test new hypotheses by generating controlled replicates for empirical or theoretical studies. Like classic subnetwork models before, our multiplex model, and most importantly, the idea behind our model – that the non-independence of traits generating different types of interactions could lead to predictable features of the structure and stability of multiplex networks^53^ – can be a useful tool for developing a more holistic understanding of complex natural systems.

## Acknowledgments

This research was supported in part through computational resources and services provided by Advanced Research Computing at the University of Michigan, Ann Arbor. We thank George Kling for comments on the manuscript. This material is based upon work supported by the NSF Postdoctoral Research Fellowships in Biology Program under Grant No. 2410512.

## Author contributions

Conceptualization: KRSH, ET, FSV

Methodology: KRSH

Analysis: KRSH

Software: KRSH

Writing – original draft: KRSH

Writing – review & editing: KRSH, ET, FSV

## Data and materials availability

All code, including sample networks and visualizations, is available by request.

## Supplementary Materials

Materials and Methods

Supplementary Text

Supplementary Methods

Figs. S1 to S6

## Materials and Methods

### Methods

#### 1. The multiplex network model

Multiplex ecological networks are a type of multilayer network in which the same set of species can participate in multiple interaction types (formally, a “node-aligned multilayer network” in which nodes – representing species – are perfectly coupled across layers – representing different interaction types)^16,19^. Here, we refer to the different interaction types as “subnetworks” (rather than layers)^18^, focusing our study on how the interconnection between subnetworks into multiplex networks emerges from shared species and trait structure. We simulate these networks by first generating communities of species with multivariate trait distributions, and then applying trait-based rules to determine interactions within each subnetwork.

##### 1.1. Species traits

We begin by modeling each species in a regional pool as having a vector of *n* continuous trait values, with each trait (*t*_*x*_) corresponding to one interaction type. These traits may be measurable or abstract, i.e., representing a single ecological trait, a composite of multiple traits, or even a latent structuring variable^4^. In natural communities, traits (*t*_*x*_, *t*_*y*_) are not independent – biological constraints such as allometry, genetic correlations, ecological trade-offs, and shared evolutionary history often cause trait values to covary. To model this dependence structure, we use a copula, a multivariate distribution that separates marginal trait distributions from their dependence structure.

According to Sklar’s theorem^89^, any multivariate joint distribution can be decomposed into the univariate marginal distributions for each trait, and a copula function that describes how these traits covary. The copula^90^ captures the rank correlations between traits and allows for arbitrary continuous marginal distributions for each trait (e.g., uniform, normal, log-normal). We use a Gaussian copula to accommodate *n* trait axes simultaneously; its untransformed marginal distributions are uniform on the unit interval *U*(0,1). Here, we use these uniform marginal distributions for consistency with classic subnetwork models, and without loss of generality, but copulas support any marginal trait distributions (including different distributions for different traits) and can even be flexibly fit to empirical data.

We define a symmetric, positive semi-definite correlation matrix ***T***_*n*×*n*_ with entries *τ*_*xy*_ representing the rank correlations between trait axes *t*_*x*_ and *t*_*y*_, with *τ*_*xx*_ = 1, and *τ*_*xy*_ = *τ*_*yx*_ to represent the commutativity of trait correlations. Because ***T*** must be positive semi-definite, note that not all trait correlation combinations are feasible: if *τ*_*xy*_, *τ*_*xz*_ are strongly positive, trade-offs among dimensions dictate that *τ*_*yz*_ must be strongly negative. Then, we randomly sample the copula *S*_*R*_ times to create a regional species pool of *S*_*R*_ species, where each species *i* has a set of *n* trait values 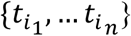 constrained by the specified trait correlations. At the community scale (the joint community composition), each trait is uniformly distributed and respects the specified rank correlations.

##### 1.2. Simulating multiplex networks

Each multiplex networks consists of *n* subnetworks corresponding to a single type of interaction (e.g., predator-prey, parasitoid-host, pollinator-plant). For each subnetwork *x* ∈ {1, …, *n*}, we specify the number of animal (*A*_x_) and plant (*P*_*x*_) species, such that *S*_*x*_ = *A*_x_ + *P*_*x*_, the number of target interactions (links, *L*_x_), and an interaction rule that determines whether two species interact based on their trait values (see *Interaction rules*, below).

We simulate each subnetwork by randomly sampling *S*_*x*_ local species from the regional pool of *S*_*R*_ species and applying the interaction rule. Following classic subnetwork models, we accept the subnetwork if all *S*_*x*_ species have at least one link (no isolated nodes), and the realized *L*_x_ is within 3% of the target links. We allow up to 1,000 tries to simulate each subnetwork, resampling *S*_*x*_ species from the regional pool each try. We repeat this process for each subnetwork; the order in which subnetworks are simulated does not appear to affect the structural properties of the subnetworks or the multiplex network.

Because species are sampled from a shared regional pool with a joint trait structure, the overlap of species identities into subnetworks (i.e., species that participate in multiple interaction types) naturally emerges from the model, and is ultimately governed by trait correlations and sampling. We may choose to reject a simulated multiplex network if it does not fulfill additional criteria suited to its ecological interpretation, such as that every local consumer has at least one resource species (i.e., in any subnetwork). These criteria act as ecological filters, shaping the local community composition and species’ functional roles therein, because only subsets of the regional pool that satisfy the interaction rules, under the constrained trait structure, are included. In particular, if all *S*_*x*_ < *S*_*R*_, the unique number of species *S* in the local multiplex network can be less than the regional richness *S*_*R*_ unless a target *S* is used as an additional filter. In the main text experiments, we chose parameters that led to *S = S*_*R*_ with no additional filters required.

###### 1.2.1. Interaction rules

We modeled species interactions within each subnetwork using rules derived from classic generating models for ecological networks. These rules define whether a pair of species interact given their trait values and a target subnetwork connectance *C*_*x*_, defined as the fraction of all potential links in subnetwork *x* that are realized.

Following (*20*), we apply these rules to construct either *unipartite* subnetworks (e.g., food webs/predator-prey) or *bipartite* subnetworks (e.g., herbivore-plant, pollinator-plant). *Unipartite*: All species *S*_*x*_ can potentially interact with all other species. Connectance is 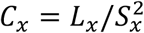. *Bipartite*: Species are divided into two groups (e.g., animals and plants), 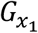 and 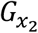. All species in 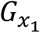 can potentially interact with all species in 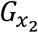, but no within-group interactions are permitted. Connectance is 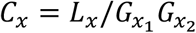, which is *C*_*x*_ = *L*_x_/*A*_x_*P*_*x*_ when the groups are defined to be animals and plants.

We use the following rules: *Threshold rule*: Species *i* with trait value 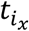can interact with species *j* with trait value 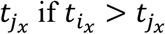. There is a probability *p*_x_ that each of these potential interactions is realized. For unipartite subnetworks, *p*_x_ = 2C_*x*_*S*_*x*_/(S_*x*_ ™ 1) (aka, the Original Cascade Model^58^. For bipartite subnetworks^68^, *p*_x_ = *L*_x_/*N*_x_ where *N*_x_ is the total number of trait-compatible pairs. This rule hypothesizes a hierarchy in which species can only interact with partners of a lower trait value (e.g., larger predators eating smaller prey). *Interval rule*: Species *i* with trait value 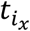 interacts with all partners *j* whose trait 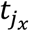 falls within a range (interval) of the trait axis 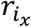, centered around a given value 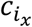, such that 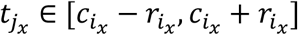. The center of the interval 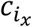 is sampled from a uniform distribution with 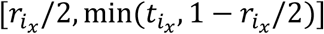 (see^8^) and the range 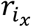 is defined as 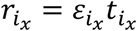, where 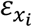 is sampled from a beta distribution with parameters *α*_x_ = 1, *β*_x_ = (1/2C_*x*_) ™ 1. In unipartite subnetworks, all species are assigned interaction ranges and centers (aka, the Original Niche Model^2^). In bipartite subnetworks^68^, only species in one group 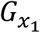 (e.g., animals) are assigned interaction ranges and centers. This rule hypothesizes trait matching/filtering (e.g., pollinators visiting flowers within a size range) combined with a looser version of the threshold hierarchy in which species tend to interact only with partners of a lower trait value. *Random rule*^58^: Species interact randomly, regardless of trait values. In unipartite subnetworks, any species can potentially interact with any other species. In bipartite subnetworks, species in one group 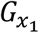 can interact with any species in the other group 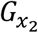. The probability that each of these potential interactions is realized is *p*_x_ = *C*_*x*_.This rule hypothesizes that species’ interactions are independent of their trait values.

#### 2. Stability of multiplex communities

##### 2.1. Multiplex networks

We investigated the effect of trait composition on the local stability of multiplex networks using a classic community matrix approach. Specifically, we studied multiplex networks of three types of co-occurring interactions that are fundamental to natural systems (citation): (“*P*”) a unipartite predator-prey subnetwork with antagonistic (+/–) effects (i.e., predators benefit and prey are harmed from the interaction), (“*H*”) a bipartite herbivore-plant subnetwork with antagonistic (+/–) effects, and (“*M*”) a bipartite animal-plant mutualist subnetwork with mutually beneficial (+/+) effects. In this section, we notate plant richness as *R* to distinguish it from index for the predator-prey subnetwork and use ***x***^*^ for equilibrium abundance.

For each feasible combination of pairwise trait correlations between subnetwork trait axes *τ*_*HP*_, *τ*_*HM*_, *τ*_*MP*_ = {™1, ™0.9, …, +0.9, +1}, we simulated a set of *N* = 300 multiplex network replicates (specifically, we simulated *N* = 100 multiplex networks by iteratively generating subnetworks in each of the following orderings: {*H, M, P*}, {*H, P, M*}, {*P, H, M*}; simulation order does not affect the structural properties). Trait combinations are feasible if they create a positive semi-definite correlation matrix. Infeasibly arises when the strong trait correlation(s) in one or more dimensions constrains the possible values of the other trait correlations. We constructed the multiplex networks using threshold rules for each subnetwork, with richnesses and connectances fixed as follows: *S* = *A* + *R* = 60 = 40 + 20, *S*_*P*_ = *A*_P_ = 40, *S*_*H*_ = *A*_H_ + *R*_H_ = 20 + 20, *S*_*M*_ = *A*_M_ + *R*_M_ = 20 + 20, *C*_*P*_ = 0.1 (≈ 160 links), *C*_*H*_ = *C*_*M*_ = 0.2 (≈80 links each).

##### 2.2. Community matrices

The topology of each multiplex network (*who interacts with whom*) can be represented in an *S* × *S* × *n* tensor ***Z***, where 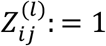 if consumer (*i*) feeds on resources (*j*) in subnetwork (*l* ∈ {*H, M, P*}), or 0 otherwise. For each network, we analyzed the stability of its *S* × *S* community matrix (***A***), with entries *a*_i*j*_ parameterized according to the following ecological assumptions:

i. *Equilibrium and dynamics:* Communities follow generalized Lotka-Volterra dynamics *d*x_*i*_/*d*t = (*r*_i_ + ∑_*j*_ *a*_i*j*_x_*j*_ ™ *a*_i*i*_*x*_*i*_)x_*i*_, with additive interspecific interaction strengths (*a*_i*j*_) across subnetworks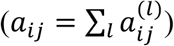. Equilibrium abundances are arbitrarily set to the feasible value of 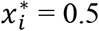, intraspecific interaction strengths are arbitrarily set to *a*_i*i*_ = 1, and *r*_i_’s are chosen so that *d*x_*i*_/*d*t = 0 at ***x***^∗^.
ii. *Energetic constraints:* Consumers (*i*) and their resources (*j*) exert reciprocal effects on each other 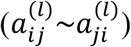, but consumers exert stronger effects on resources than the benefits they receive 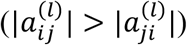 to reflect assimilation inefficiencies and energetic costs. Specifically, we chose 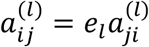, with *e*_*l*_ = +1/6 for mutualistic interactions (*l* = *M*) and ™1/6 for antagonistic interactions (*l* ∈ {*H, P*}).
iii. *Generalist-specialist trade-off:* consumers distribute total foraging effort equally among their resources, whether prey, plant, or mutualist, such that more specialized consumers have stronger effects on their resources than generalists: 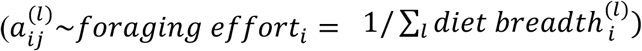. Specifically, 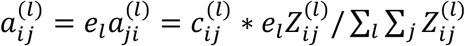, where 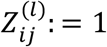 if consumer (*i*) feeds on resources (*j*) in subnetwork (*l*), or 0 otherwise. We arbitrarily set the constant 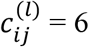. Small amounts of stochasticity in any of the arbitrarily chosen values do not qualitatively affect our results.

These assumptions are similar to those of previous authors studying the stability of complex communities^46–48,63,86^. A community is said to be locally stable if, following a small perturbation, it returns to equilibrium ***x***^∗^, that is, the real part of the dominant eigenvalue of the Jacobian of the community matrix is negative, 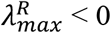. Resilience is defined by the rate of return to equilibrium following a small perturbation, quantified as 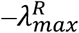; by using this convention, we can say that as 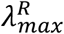 becomes more negative, communities become more resilient.

##### 2.3. Drivers of community stability

###### 2.3.1. Complexity

Classical work by Robert May^50^ argued that randomly generated community matrices have a very high probability of stability when 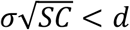, where *S* is the diversity of the community, ™*d* is the coefficient of self-limitation (i.e., diagonal entries, *a*_i*i*_) for all species, and interspecific interaction strengths (i.e., values of off-diagonal entries) are sampled from a standard normal distribution *a*_i*j*_~*N*(0, *σ*^2^) with probability *C* or are set to 0 otherwise; *C* therefore sets the connectance of the community (i.e., is the fraction of nonzero off-diagonal entries, rather than the fraction of realized links in a network as used for topological studies). Conversely, if 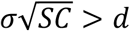, that is, at high levels of complexity (diversity, complexity, or variance in interaction strengths), randomly generated communities have a very high probability of instability. Subsequent authors^46–48,51,84,91^ have generalized this analytical expression to account for nonzero mean interspecific interaction strengths (i.e., *a*_i*j*_~*N*(*µ, σ*^2^), where *µ* is the mean of the absolute value of nonzero off-diagonal entries), as well as forms of nonrandom structure common in natural ecological communities, such as reciprocal / pairwise interactions potentially with correlated effects (i.e., *a*_i*j*_~*a*_*j*i_, with Pearson correlation among reciprocal interaction strengths *ρ*), arbitrary equilibrium distributions ***x***^∗^, or specified sign structures, like fractions of positive or negative interactions (e.g., *a*_i*j*_~ + |*N*(0, *σ*^2^)| with probability *p*C or *a*_i*j*_~ ™ |*N*(0, *σ*^2^)| with probability (1 ™ *p*)C; a similar approach is used to specify fractions of randomly assigned mutualistic or antagonistic interactions).

In our work, diversity of the community (*S*) and the number and signs of distinct nonzero interactions is fixed, as are the broad topological properties of each individual subnetwork. However, multiplex network structure varies with underlying community trait structure, with consequences for the distribution of interaction strengths (*µ, σ*^2^, *ρ*), effective connectance (*C*), and therefore the stability of the community. We report our results in terms of May’s classic complexity metric, defined as 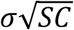. We show the individual effects of (*µ, σ*^2^, *ρ, C*) on stability in **Fig. S6**.

###### 2.3.2. Properties independent of complexity

Stability is also driven by localized concentrations of mutualistic interactions within community, which we report in terms of the maximum total mutualistic foraging effort on an individual plant species (*k*): 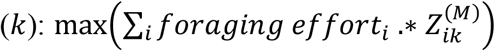, where 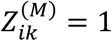 if *i* is a mutualistic consumer of *k* or 0 otherwise. Note that high values of mutualistic foraging effort are driven by hyperspecialist mutualist consumers, whose *foraging effor*t_*i*_ = 1. To check that concentrated strong mutualistic interactions affect stability independent of complexity, we chose a subset of 240 of the original 4,787 treatments that span the space of feasible trait correlations.

For each replicate community matrix (***A***) in each treatment, we created 100 matching randomized versions (**Ã**) with exactly matched complexity and reciprocal sign structure. Note that the community matrix ***A*** has entries 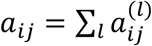, representing the effect a consumer *i* has on resource *j*, and reciprocal entries 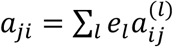 representing the effect of the resource on the consumer; therefore, the consumers’ effects are confined to the lower triangular part of ***A*** while the resources’ effects are mirrored in the upper triangular part. So, to create a randomized version of ***A***, we randomly re-assigned each 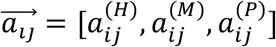 to another entry in the lower triangular part of **Ã** without replacement and then used these entries to calculate the reciprocal effects of the resource in the upper triangular portion of **Ã**. Note that shuffled matrices**Ã**. have identical distribution of interaction strengths (*µ, σ*^2^, *ρ*), effective connectance (*C*), and diversity (*S*) as the original matrix ***A***. From there, we proceeded to evaluate the maximum total mutualistic foraging effort of **Ã** and its resilience. For each shuffled matrix **Ã**, we calculated the difference between the shuffled value and the original value for the community matrices from the multiplex treatments. We report the mean resilience of all the shuffled matrices for all the replicates in a treatment, as well as the mean differences in resilience and maximum total mutualistic foraging effort.

###### Materials

Network simulations and topological analyses were performed in MATLAB R2024b (Mathworks) and bipartite network analyses (modularity and nestedness) in the supplement were calculated using the BiMat package^92^. All code, including sample networks and visualizations, is available by request.

