## Supplementary for "Correlated traits generate the structure and stability of multiplex ecological networks"

### Supplementary Text

#### An empirical example

Classic generating models use a single trait axis and simple interaction rules to generate ‘subnetworks’ of a single interaction type. Our model is a generalization of these previous models using multiple, interdependent trait axes to generate ‘multiplex networks’ of multiple interaction types. As such, it can be used to explicitly encode and test hypotheses (as interaction rules) about how species traits jointly determine interactions within and across different subnetworks, while still replicating the known features of those subnetworks embedded in multiplex networks. In our construction, we consider the simplest case – that the underlying trait axes are correlated but that interaction rules are applied independently to generate each subnetwork.

To illustrate this, we provide an empirical example of conservation interest. In habitats that are depauperate of other resources, some bird species forage on both nectar and fruits or seeds, acting as both pollinators and seed dispersers (“double mutualists”)<sup>78,80</sup>. Because traits underlying these interactions are positively correlated across all bird species ($\tau_{obs} \approx 0.74$ for beak widths and lengths)<sup>4,81,93</sup>, we expect that the bird species that are perhaps the most important pollinators in these environments – i.e., generalists, pollinating the most plant species – are doubly important because they are also generalist seed dispersers – dispersing the most plant species. Specifically, given the observed correlation among bird traits, the number of plant and bird species, and the connectance of each subnetwork (see Supplementary Methods), our model reproduces key properties of an empirical multiplex network of pollination and seed dispersal interactions from the Galapagos Islands archipelago<sup>79</sup> (**Fig. S1A**).

We simulated multiplex networks by assuming that the pollination and seed dispersal subnetworks were generated by trait axes correlated with coefficient $\tau_{xy} = \{-1, -0.9, \dots, +1\}$. When $\tau_{xy} = 0$, the traits that underly pollination and seed dispersal interactions are independent of each other; when $\tau_{xy} = +1$, they are perfectly correlated. To generate subnetworks, we considered three simple rules, widely used in classic subnetwork models: (1) the *threshold rule*, where a species can potentially interact with any partner below a threshold trait value<sup>58,68</sup>, (2) the *interval rule*, where a species interacts with all partners within a trait range (i.e., an interval)<sup>2,68</sup>, and (3) a *random rule*, where species can potentially interact with any partner, independent of trait values. The threshold rule has been shown to reproduce properties of mutualistic subnetworks in previous studies<sup>68</sup>.

Our simulation process, which randomly samples species from each trait axis to generate its corresponding subnetwork, led to a greater overlap in the plant species composition of the subnetworks than observed in the empirical network (**Fig. S2**). That is, a random (neutral)

sampling of the plant community led to significantly more species with both bird-pollination and bird-seed dispersal interactions than is observed empirically, demonstrating a nonrandom partitioning of interaction types among the species in this system. In the subsequent analyses, we used simulated multiplex networks with composition overlaps matching the empirical multiplex network (see Supplementary Methods), but this does not qualitatively affect our results.

Treatments with multiplex networks simulated from threshold rules for both the pollination and seed dispersal interactions did not significantly differ from the empirical observation in characteristic properties of mutualistic subnetworks, including their nestedness and modularity, as quantified by normalized model errors (**Fig. S1B**), and the distributions of species' degrees (i.e., number of partners within a subnetwork) as assessed by two-tailed Kolmogorov-Smirnov tests (**Fig. S1C**).

Beyond subnetwork structure, our simulated treatments also accurately reproduced emergent properties of the empirical multiplex network (**Fig. S1B-E**), including the nestedness and modularity of the aggregated bird-plant mutualism network (i.e., with distinctions between interaction types removed), the distribution of species' multiplex degree (i.e., total number of partners across subnetworks), and the distributions of species' participation coefficients (i.e., evenness of partners across subnetworks<sup>45</sup>). In particular, the high overlap between the most generalist pollinators and seed dispersers expected given $\tau_{obs}$, as well as the overlaps between other functional roles, closely match the overlaps in the empirical network (**Fig. S3**). Overall, these results highlight that our model is able to reproduce key structural properties of empirical multiplex networks, extending and integrating existing subnetwork models into a unified multiplex framework.

Interestingly, however, overlaps between the functional roles of plants across subnetworks were reproduced inconsistently across trait correlations and interaction rules (**Fig. S4**), even after correcting for the nonrandom partitioning observed in the empirical network (**Fig. S2**). This may suggest that additional processes may underly the structuring of plants as double mutualists in this system. For example, indeterminate growth, dynamic tissue allocations, and interference among plant visitors may create a more complex relationship between plant pollination and seed dispersal traits than can be described by a simple rank correlation coefficient. Deviations from the patterns predicted by our model can be used to stimulate new hypotheses for assembly processes in natural systems.

### Supplementary Methods

#### Analyzing network structure

For each of our simulated multiplex networks, we calculated *multiplex properties*, which characterize the interconnection between the $n$ subnetworks of different interaction types; *monoplex properties*, which characterize overall structural features without regard to different interaction types; and *subnetwork properties*, which characterize the structural features of each individual subnetwork.

#### Monoplex network structure

In a monoplex network, link presence is retained but interaction types are ignored (i.e., a ‘flattened’ or layer-aggregated version of a multilayer network). This allows us, for example, to compare emergent multiplex food webs (constructed from subnetworks generated with underlying trait constraints) differ from traditional food webs built from classic subnetwork models that treat all interactions equivalently.

We calculated statistics for the monoplex network version of (i) the multiplex network composed of $n$ subnetworks and (ii) the (multiplex) networks composed of each of the unique sets of $\{1, 2, \dots, n-1\}$ subnetworks that compose the multiplex network. We also calculate overlaps between species’ functional roles within the (monoplex) network (see below). When analyzing the monoplex network, we excluded species that do not participate in any of the included subnetworks. The monoplex of a single subnetwork $x$ thus has exactly $S_x$ species.

#### Multiplex network structure

To describe how subnetworks are interconnected into multiplex networks, we developed a new suite of statistics based on the overlap of species across subnetworks into multiple functional roles. We define these functional roles in a series of categories for each species in the community: inclusion (presence or absence in a subnetwork), animal or plant, consumer or resource in each subnetwork, extreme traits (top or bottom third of ranked trait values in each subnetwork), and extreme degree (top or bottom third of ranked degrees in each subnetwork). Degree ($k_{i_x}$) is the number of links a species $i$ has in subnetwork $x$. This can be interpreted as diet breadth, (mutualistic) generality, or vulnerability to consumers, depending on the interaction type and directionality of the links. By focusing on species in the top or bottom third of trait or degree rankings, we can characterize overlaps between ecologically distinct and intuitive roles (e.g., the shortest beaked and the most specialist (lowest out-degree) pollinators in a community), and track how these roles translate across interaction types into multiplex functional roles.

To quantify the overlap in identify between sets of species with different roles (both within and across subnetworks), we use the Jaccard index<sup>94</sup> (also called the Tanimoto similarity index<sup>95</sup>):

$$J(f_{i_x}, f_{j_y}) = \frac{|f_{i_x} \cap f_{j_y}|}{|f_{i_x} \cup f_{j_y}|} \quad \text{Eqn. S1}$$

which describes the number of species intersecting between $f_{i_x}$ and $f_{j_y}$, the sets of species with functional role $i$ in subnetwork $x$ and role $j$ subnetwork $y$, respectively, relative to the total number of unique species (the union) in $f_{i_x}$ and $f_{j_y}$. The expected value for the Jaccard index can be calculated analytically as:

$$\mathbb{E}[J(f_{i_x}, f_{j_y})] = \frac{\mathbb{P}(f_{i_x} \cap f_{j_y})}{\mathbb{P}(f_{i_x}) + \mathbb{P}(f_{j_y}) - \mathbb{P}(f_{i_x} \cap f_{j_y})} \quad \text{Eqn. S2}$$

where $\mathbb{P}(f_{ix})$ is the probability that any given species is a member of $f_{ix}$ and $\mathbb{P}(f_{ix} \cap f_{jy})$ is the joint probability that species are a member of both functional roles. If the two functional roles are independent (i.e., traits are uncorrelated $\tau_{xy} = 0$, or do not determine degree [random rule]). Eqn. S2 reduces to:

$$\mathbb{E}[J(f_{xi}, f_{yj})] = \frac{\mathbb{P}(f_{ix})\mathbb{P}(f_{jy})}{\mathbb{P}(f_{ix}) + \mathbb{P}(f_{jy}) - \mathbb{P}(f_{ix})\mathbb{P}(f_{jy})} \quad \text{Eqn. S3}$$

This provides a null expectation; deviations from independence show how ecological structure affects the overlap of functional roles (**Figs. S1-S4**). Eqns. S1-S3 generalize to $n > 2$ subnetworks.

To complement previous work<sup>16,18,24</sup> we calculated the realized correlation between species' traits and species' degrees in each pair of subnetworks (only including species that are present in both subnetworks), species' total degree across subnetworks ($k_i = \sum_x^n k_{ix}$), and the distribution of species' participation coefficients<sup>45</sup> across subnetworks:

$$q_i = 1 - \sum_x^n \left( \frac{k_{ix}}{k_i} \right)^2 \quad \text{Eqn. S4}$$

The participation coefficient characterizes the evenness of species' degree across subnetworks, where a species with interactions in only one subnetwork has $q_i = 0$ while a species with interactions equally distributed across subnetworks has a perfect $q_i = 1$.

### Experimental design

#### Analyzing an empirical network

We used our model to analyze an empirical multiplex network ($S = 129, A = 21, P = 108$) of bird pollination ($S_x = 66, A_x = 19, P_x = 47, C_x = 0.148$) and seed dispersal ($S_y = 100, A_y = 19, P_y = 81, C_y = 0.242$) from the Galapagos Islands archipelago (50). To match the assumptions of classic subnetwork models, we began by aggregating the taxonomic species in the empirical network into 'trophic species', wherein species with the same set of partners in both subnetworks (i.e., identical multiplex edge sets) are lumped into a single node. Because of the nestedness of mutualist subnetworks, this tended to group specialists into singular trophic species, increasing the connectance ($C_x = 0.154, C_y = 0.292$) and reducing the richness in the subnetworks ($P_x = 44, P_y = 62$) and in the overall multiplex network ($S = 107, A = 21, P = 86$).

We simulated multiplex network treatments with matched richnesses and connectances to the empirical network for the full factorial combination of: interaction rules to generate the pollination and seed dispersal subnetworks $x, y \in \{\text{threshold (T)}, \text{interval (I)}\}$; and correlation coefficients $\tau_{xy} \in \{-1, -0.9, \dots, +0.9, +1\}$ between the pollination and seed dispersal trait axes. We additionally experimented with using separate copulas for plant versus animal traits by specifying correlation coefficients $\tau_{xy}^P, \tau_{xy}^A$, respectively. As a control, we also considered treatments generated from *random* (R) rules for both subnetworks. Together, this yielded 5 rule-sets $\times 21 \tau_{xy}^P \times 21 \tau_{xy}^A = 2205$ treatments, each composed of $N = 100$ multiplex network replications.

In our simulation process, species are randomly sampled into each subnetwork, and multiplex structure emerges from the overlap of species into multiple subnetworks (e.g., the overlapping plant species are those that are both pollinated and dispersed by birds in the Galapagos). Inspecting our simulated networks, we see a range in the level of overlap among plants (**Fig. S2**), but across replicates there are generally fewer overlapping plant species than would be expected from neutrally (randomly) sampling the species pool. That is, overlap

$J(p_x, p_y)_{oemp} = 0.233 < \text{expected neutral overlap } \mathbb{E}[J(p_x, p_y)] = 0.427$ from Eqn. S2, where
$p_x = P_x/P = 44/86$, $p_y = P_y/P = 62/86$. This demonstrates a nonrandom partitioning of
interaction types among the Galapagos plants species. To account for this, we also experiment
with using a larger regional plant pool $P_R > P$ such that the emergent plant overlap in our
simulations matches the empirical value. Specifically, we calculated $P_R = 136$ by solving Eqn.
S2 using $\mathbb{E}[J(p_x, p_y)] = J(p_x, p_y)_{emp} = 0.233$, $p_x = P_x/P_R = 44/P_R$, and $p_y = P_y/P_R =$
$62/P_R$. Then, we re-simulated our network treatments and filtered the multiplex network
replicates until we had $N = 100$ replicates with exactly the total empirical plant richness of $P =$
86 for each treatment. These results are shown in **Figs. S3-4**, but do not qualitatively differ from
results without this correction.

Plant-animal interaction networks are commonly characterized<sup>56</sup> by their modularity (in
which a subset of nodes more frequently connected to each other than to others) and nestedness
(in which specialists interact with subsets of the partners of generalists). To compare empirical
and simulated subnetworks, we calculated the standardized modularity and nestedness for each
subnetwork, as well as for the monoplex version of the multiplex bipartite network. In all cases,
for the focal network, we calculated modularity using the leading eigenvector method<sup>96</sup> and
nestedness using the NODF statistic<sup>97</sup> as implemented in the BiMat package<sup>92</sup>. Then we
standardized these values to a set of 100 null networks with matching row and column degrees
(the “FIXED” null model) as $(\text{focal} - \text{mean}(\text{null set})) / \text{mean}(\text{null set})$. Additionally, we recorded
the degree distributions for all species, animal consumers (birds), and plants separately in each
subnetwork, and summed as double mutualists (multiplex degree) in the multiplex network.
Finally, we recorded overlaps of functional roles and the distribution of participation coefficients
for all species and separately for birds and plants. We used two-tailed Kolmogorov-Smirnov tests
to test the null hypothesis that the distributions in simulated and empirical networks were
sampled from the same underlying distribution.

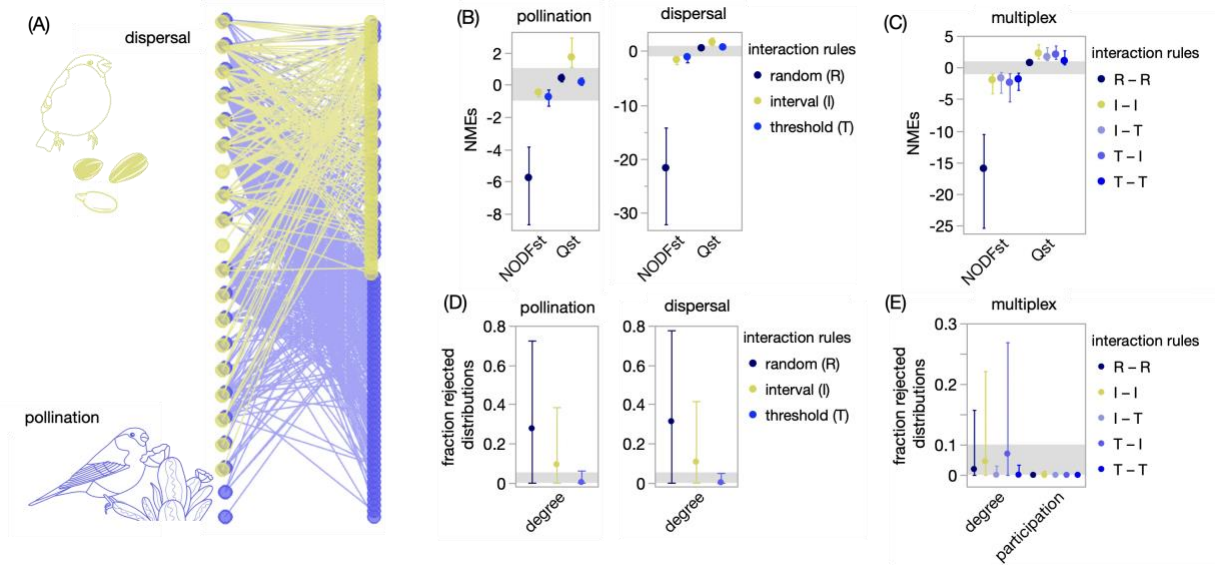

**Fig. S1. Multiplex and subnetwork properties of an empirical pollination–seed dispersal network are well-predicted by the multiplex model using threshold rules.** (A) An empirical multiplex network of bird-pollination and -seed dispersal interactions from the Galapagos Islands (50). Vertically aligned nodes indicate that the same bird (purple) or plant (yellow) species engages in both the (i) pollination and (ii) seed dispersal subnetworks. (B–C) Normalized model errors (NMEs) are the normalized differences between the empirical observation and the median values for treatments of 100 multiplex networks simulated according to each combination of trait correlations and interaction rules (colors): interval (I), threshold (T), random (R); $N = 2205$ total treatments. We show the mean (circles) and range (error bars) of NMEs for the nestedness and bipartite modularity of (B) simulated subnetworks, and (C) their overall multiplex networks. $NMEs < 0$ ($> 0$) indicate that simulated networks underestimate (overestimate) an empirical property; values between $(-1, +1)$, gray bar) indicate no significant difference at the 95% confidence level. (D–E) Fractions of simulated networks for which two-tailed Kolmogorov-Smirnov tests reject the null hypothesis that properties of simulated and empirical networks were sampled from the same underlying distribution (treatments are considered to significantly differ if the null hypothesis is rejected for $\geq 5\%$ of their simulated networks). We show the mean (circles) and range (error bars) for the distribution all species' degree (number of partners) in (D) simulated subnetworks, and (E) the distribution of species' degree (total number of partners across subnetworks) and participation coefficients (evenness of partners across subnetworks) in their overall multiplex networks.

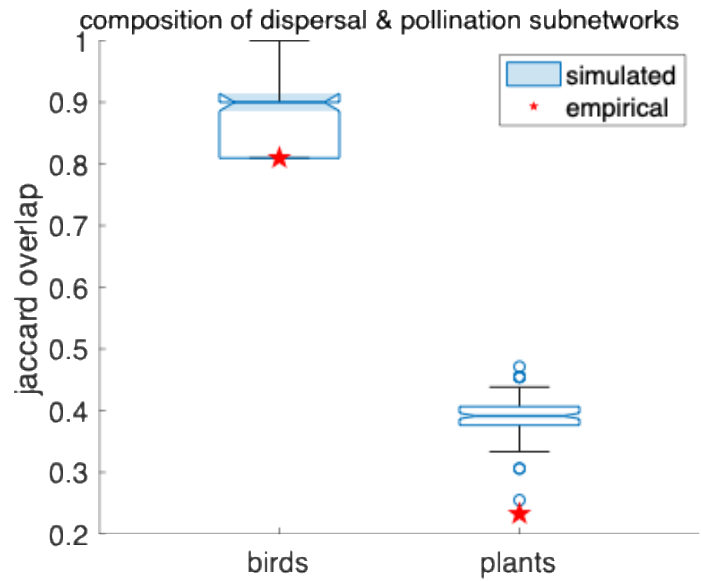

**Fig. S2. Overlap of bird and plant compositions as double mutualists into seed dispersal and pollination subnetworks.**

The overlap in bird and plant species compositions among the seed dispersal and pollination subnetworks. Randomly sampling species from each trait axis to generate its corresponding subnetwork led to a slightly greater overlap in animal species composition, and a significantly greater overlap in the plant species composition for simulated multiplex networks than observed in the empirical network. In other words, a random (neutral) sampling of the plant community led to substantially more species with both bird-pollination and bird-seed dispersal interactions than is observed empirically, demonstrating a nonrandom partitioning of interaction types among the species in this system. For the remaining analyses, we filtered out multiplex network simulations that did not match the observed overlaps (see Supplementary Methods).

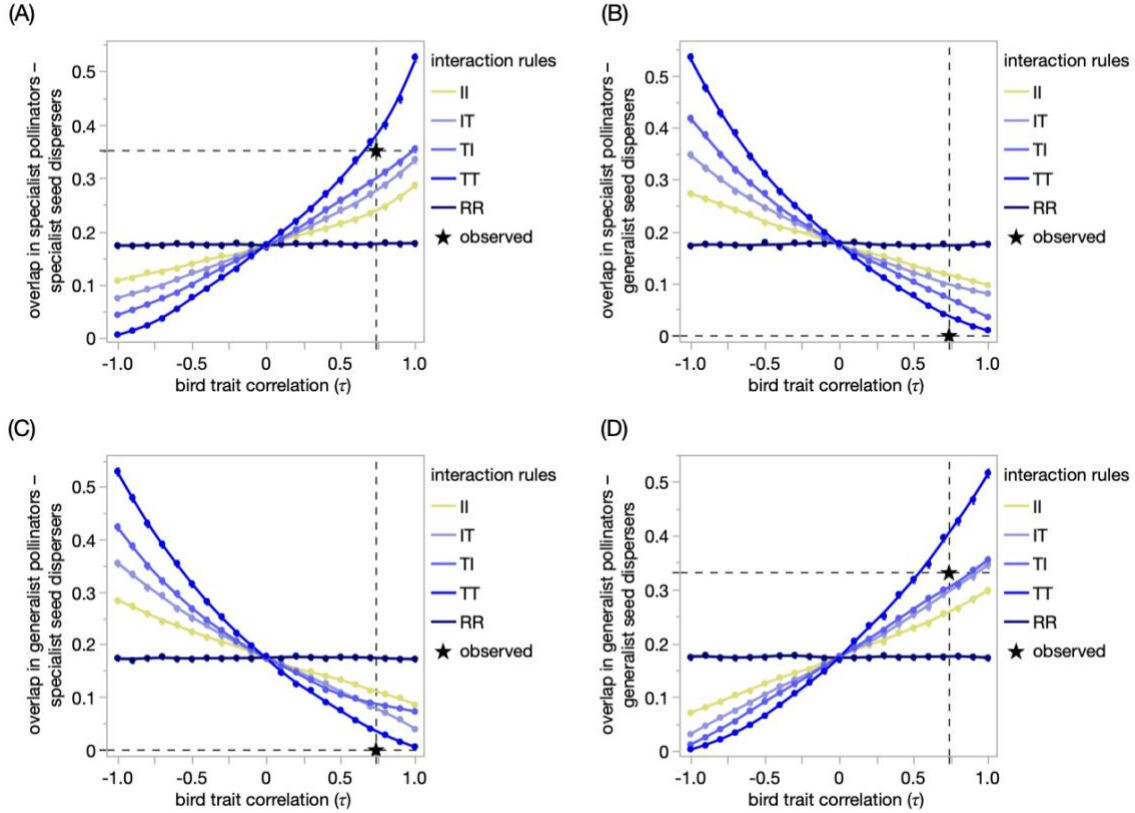

**Fig. S3. Properties of birds as “double mutualist” seed dispersers and pollinators in the Galapagos are well-predicted by the multiplex model with empirical trait correlations.**

Properties of simulated networks (colored lines) vary with the correlation between birds’ traits thought to underly pollination (beak length) and seed dispersal (beak width); the dashed vertical line marks the empirically-observed value across all bird species<sup>4,81,93</sup> ($\tau_{obs} \approx 0.74$). The dashed horizontal lines indicate the empirically observed overlap between birds’ functional roles as specialist or generalist pollinators and seed dispersers (mean Jaccard similarity from 100 bootstrapped replications). The star marks the intersection of these observed values. Empirically-observed overlap between birds’ functional roles as specialist or generalist pollinators and seed dispersers compared to overlaps in simulated multiplex networks (colored lines). Specialists are bird species in the bottom 1/3 of the consumer degree distributions (i.e., linked to the fewest resource plants), while generalists are in the top 1/3 of the consumer degree distribution (i.e., linked to the most resource plants). **(A)** Specialist pollinators and specialist seed dispersers. **(B)** Specialist pollinators and generalist seed dispersers. **(C)** Generalist pollinators and specialist seed dispersers. **(D)** Generalist pollinators and generalist seed dispersers. We show the means (points) and 95% confidence intervals (error bars) for each treatment of 100 multiplex networks simulated according to each combination of trait correlations and interaction rules (colors): interval (I), threshold (T), random (R); $N = 2205$ total treatments.

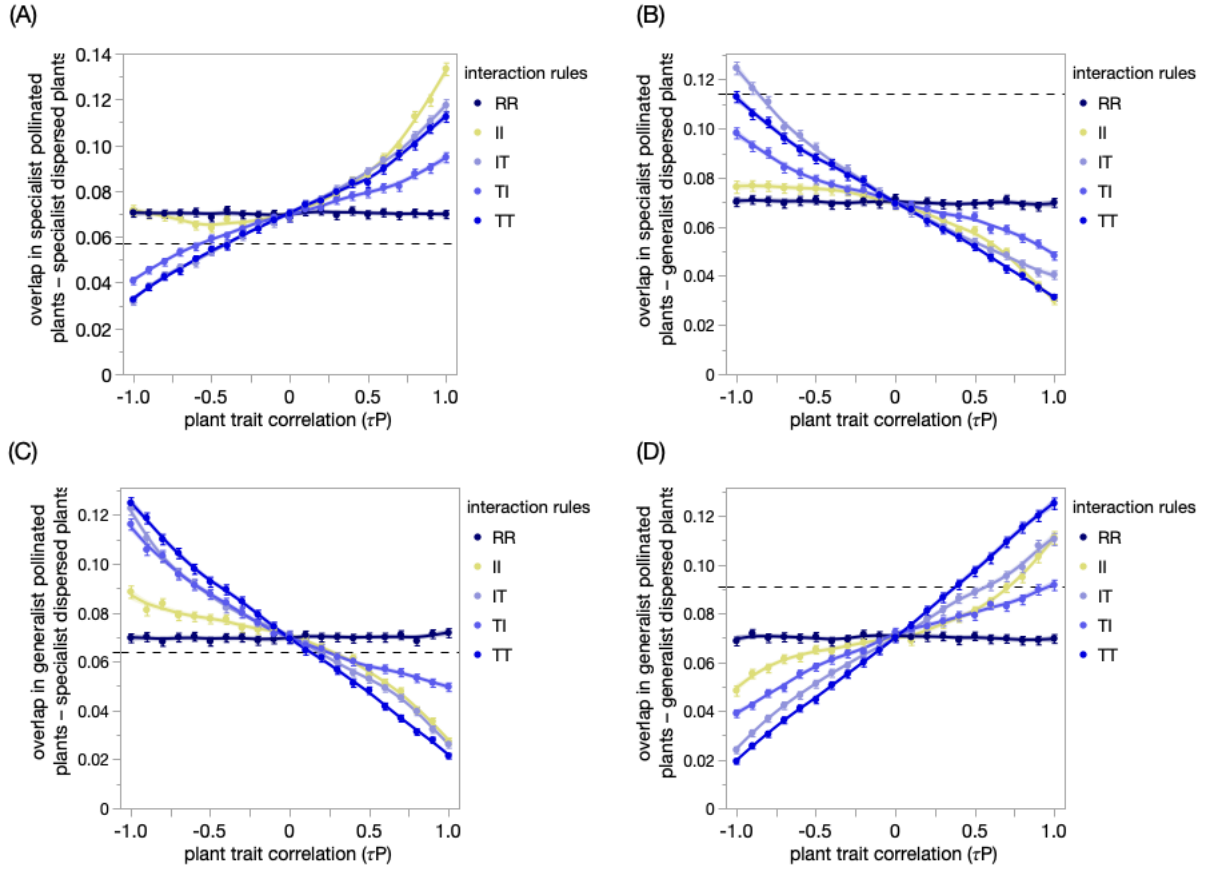

**Fig. S4. Properties of plants as “double mutualists” of pollinating and seed dispersing birds in the Galapagos.** We compare the empirically observed overlaps (dashed horizontal lines) between each combination of generalist and specialist plant mutualists to mean values (points) and 95% confidence intervals (error bars) from each treatment of 100 multiplex networks simulated according to each combination of trait correlations and interaction rules (colors): interval (I), threshold (T), random (R); $N = 2205$ total treatments. Unlike for birds, we have no a-priori expectation for a trait correlation. All other formatting follows Fig. S2.

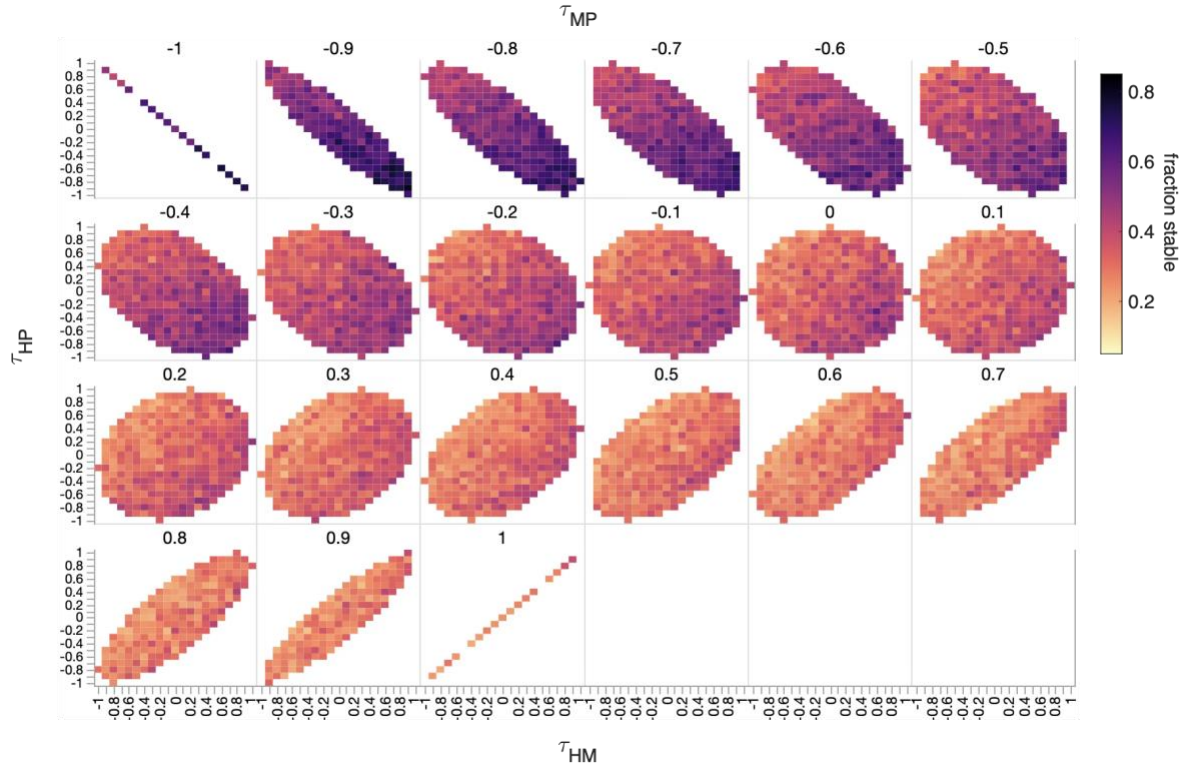

**Fig. S5.** Detailed heatmaps showing the effect of pairwise correlations ($\tau_{xy}$) among predator-prey ($P$), herbivore-plant ($H$), and animal-plant mutualist ($M$) trait axes on the community stability of multiplex networks. Multiplex networks were simulated in treatments of 300 stochastic replicates for each of the 4,787 sets of feasible trait correlations, $\vec{\tau} = [\tau_{HM}, \tau_{HP}, \tau_{MP}]$. A replicate is locally stable if its resilience ($-\lambda_{max}^R$) is positive, where $\lambda_{max}^R$ is the maximum real eigenvalue of the Jacobian of the corresponding community matrix. Heatmap cells show the fraction of locally stable replicates within a treatment. The correlation between herbivore-plant and mutualist-plant trait axes ($\tau_{HM}$) increases along the x-axis, the correlation between herbivore-plant and predator-prey trait axes ($\tau_{HP}$) increases along the y-axis, and the correlation between predator-prey and mutualist-plant trait axes ($\tau_{MP}$) increases as the panels move from left to right and wrap around; darker colors indicate greater stability.

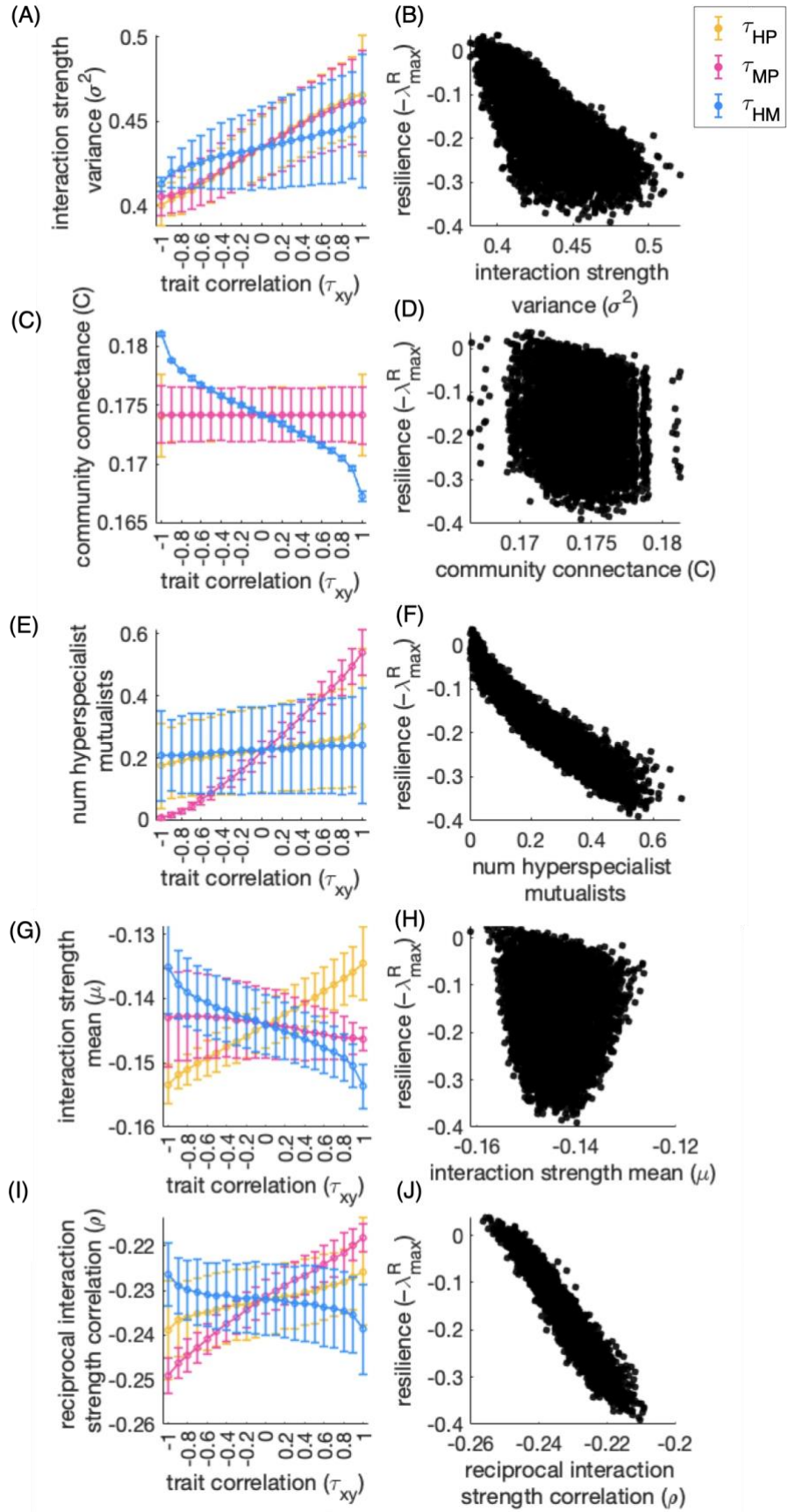

**Figure S6. Trait correlations affect resilience through different components of complexity and species' roles in the community matrix.**

Trait correlations alter emergent community complexity through the distribution and alignment of consumers' foraging effort across interaction types. Complexity ($\mu + \sigma\sqrt{SC}$) increases with the mean ($\mu$) and variance ($\sigma^2$) of interaction strengths, and the connectance of the community ($C$), while species diversity ($S$) is fixed across simulations. (A) Negative correlations among all pairwise trait combinations – predator-prey and herbivore-plant ($\tau_{HP}$, yellow), predator-prey and mutualist-plant ($\tau_{MP}$, pink) and herbivore-plant and mutualist-plant trait axes ($\tau_{HM}$, blue) – align generalist consumers in one subnetwork with specialist consumers in the other, evenly distribution foraging effort across subnetworks and decreasing interaction strength variance ($\sigma^2$); positive correlations have the opposite effect. (B) Greater $\sigma^2$ decreases resilience. (C, G) Increasing the herbivore-plant and mutualist-plant trait correlation ($\tau_{HM}$) aligns negative herbivory and positive mutualistic effects on shared plants, creating weak net effects that overall reduce mean interaction strength ($\mu$) and effective community connectance ($C$). Increasing the predator-prey and herbivore-plant trait correlation ($\tau_{HP}$) aligns generalist predators with generalist herbivores, creating weaker interactions on average. This increases mean interaction strength of the whole community, i.e., makes it less negative. (D, H) Overall reductions in $\mu$ and $C$ across our simulations are small, and have weak effects on resilience in isolation. (D) Increasing the correlation between predator and mutualist traits ($\tau_{MP}$) increases the frequency of hyperspecialist mutualists (mean number in a treatment) that exert the strongest effects in the community. (F) The frequency of hyperspecialist mutualists is associated with lower resilience. (I) Increasing the correlations among predator-prey and herbivore-plant ($\tau_{HP}$) and predator-prey and mutualist-plant ($\tau_{MP}$) tends to increase the correlation among reciprocal interaction strengths, while increasing the herbivore-plant and mutualist-plant trait correlation ($\tau_{HM}$) weakens this relationship because the additive net effects to the consumer are positive, while the net effects to the plant resource are weaker due to both positive (mutualistic) and negative antagonistic components. In (A, C, E, G, I) points and error bars show the means and standard deviations among treatments with the indicated trait correlation $\tau_{xy}$. In (B, D, F, H, J), each point ($N = 4,787$) represents the mean of one treatment ($N = 4,787$). Formatting follows Figure 3 in the main text.
